# Lagging strand encoding promotes adaptive evolution

**DOI:** 10.1101/2020.06.23.167650

**Authors:** Christopher N. Merrikh, Leonard A. Harris, Sarah Mangiameli, Houra Merrikh

## Abstract

Cells may be able to promote adaptive evolution in a gene-specific and temporally-controlled manner. Genes encoded on the lagging strand have a higher mutation rate and evolve faster than genes on the leading strand. This effect is likely driven by head-on replication-transcription conflicts, which occur when lagging strand genes are transcribed during DNA replication. We previously suggested that the ability to selectively increase mutagenesis in a subset of genes may provide an adaptive advantage for cells. However, it is also possible that this effect could be neutral or even highly deleterious. Distinguishing between these models is important because, if the adaptive model is correct, it would indicate that 1) head-on conflicts, which are generally deleterious, can also provide a benefit to cells, and 2) cells possess the remarkable ability to fine-tune adaptive evolution. Furthermore, investigating these models may address the long-standing debate regarding whether accelerated evolution through conflicts can be adaptive. To distinguish between the adaptive and neutral models, we conducted single nucleotide polymorphism (SNP) analyses on wild strains of bacteria, from divergent phyla. To test the adaptive hypothesis, we analyzed convergent mutation patterns. As a simple test of the neutral hypothesis, we performed *in silico* modeling. Our results show that convergent mutations are enriched in lagging strand genes and that these mutations are unlikely to have arisen by chance. Additionally, we observe that convergent mutation frequency has a stronger positive correlation with gene-length in lagging strand genes. This effect strongly suggests that head-on conflicts between the DNA replication and transcription machineries are a key mechanism driving the formation of convergent mutations. Together, our data indicate that head-on replication-transcription conflicts can promote adaptive evolution in a variety of bacterial species, and potentially other organisms.

## Introduction

In bacteria, the majority of genes, especially highly transcribed and essential genes, are encoded on the leading strand. This is likely due, in part, to strong negative selection against highly transcribed lagging strand alleles, which can cause severe head-on replication-transcription conflicts^1–5^. Yet after billions of years of evolution, many lagging strand genes remain^6–10^. It is likely that many head-on genes persist in this orientation because they are not transcribed during DNA replication, and therefore do not impede DNA replication (the neutral hypothesis). Together, negative selection and neutral evolution are, in theory, sufficient to explain the relatively low abundance of genes on the lagging strand^11^.

However, in 2013, we presented evidence that positive selection can also promote the retention of lagging strand alleles^7^. We found that, when transcribed, lagging strand genes mutate at a faster rate than otherwise identical leading strand genes as a result of head-on replication-transcription conflicts^7^. This effect is mirrored by the observation that, on the whole, lagging strand genes evolve at a faster rate than lagging strand genes in nature^6,7^. These findings indicate that the mutation rate of a given gene may be raised or lowered by DNA transactions (e.g. recombination) that change a gene’s coding strand, provided that it maintains an equivalent transcription profile. As such, we proposed that the mutation rate of individual genes is not a constraint, rather it can be fine-tuned through changes in gene orientation^7^. This implies that cells may gain an adaptive advantage from encoding genes that frequently come under selection on the lagging strand.

Others have offered alternative explanations for the higher mutation rate of lagging strand genes. One model suggests that differential sequence context in lagging strand genes could increase the mutation rate^12,13^. We investigated this possibility, but found no evidence supporting the model, at least in the context of natural evolution^14^. It was also proposed that the lower expression level of lagging strand genes (at least during growth in rich media) reduces the potential problems of acquiring detrimental mutations^3,11^. Such a reduction in purifying selection could allow lagging strand genes to retain more non-nonsynonymous, but otherwise non-adaptive mutations^15,16^.

It is possible to distinguish between the higher mutability hypothesis and the adaptive hypothesis. If lagging strand mutations turn out to be largely non-adaptive, it would support the mutability hypothesis. Conversely, if lagging strand mutations are more frequently beneficial than leading strand mutations, it would support the adaptive hypothesis. One way to determine if mutations are beneficial is to examine the frequency with which the same mutations arise in independent lineages. Such evolutionary convergence is a powerful indicator of positive selection^17^. Our previous SNP analysis of 5 *B. subtilis* genomes suggested that parallel and multi-hit site mutations (varieties of convergent mutations) may be more common in lagging strand genes (Fig. 1A)^7^. However, this analysis had low statistical power and as such alternative models remained viable possibilities^11^.

**Figure 1.**
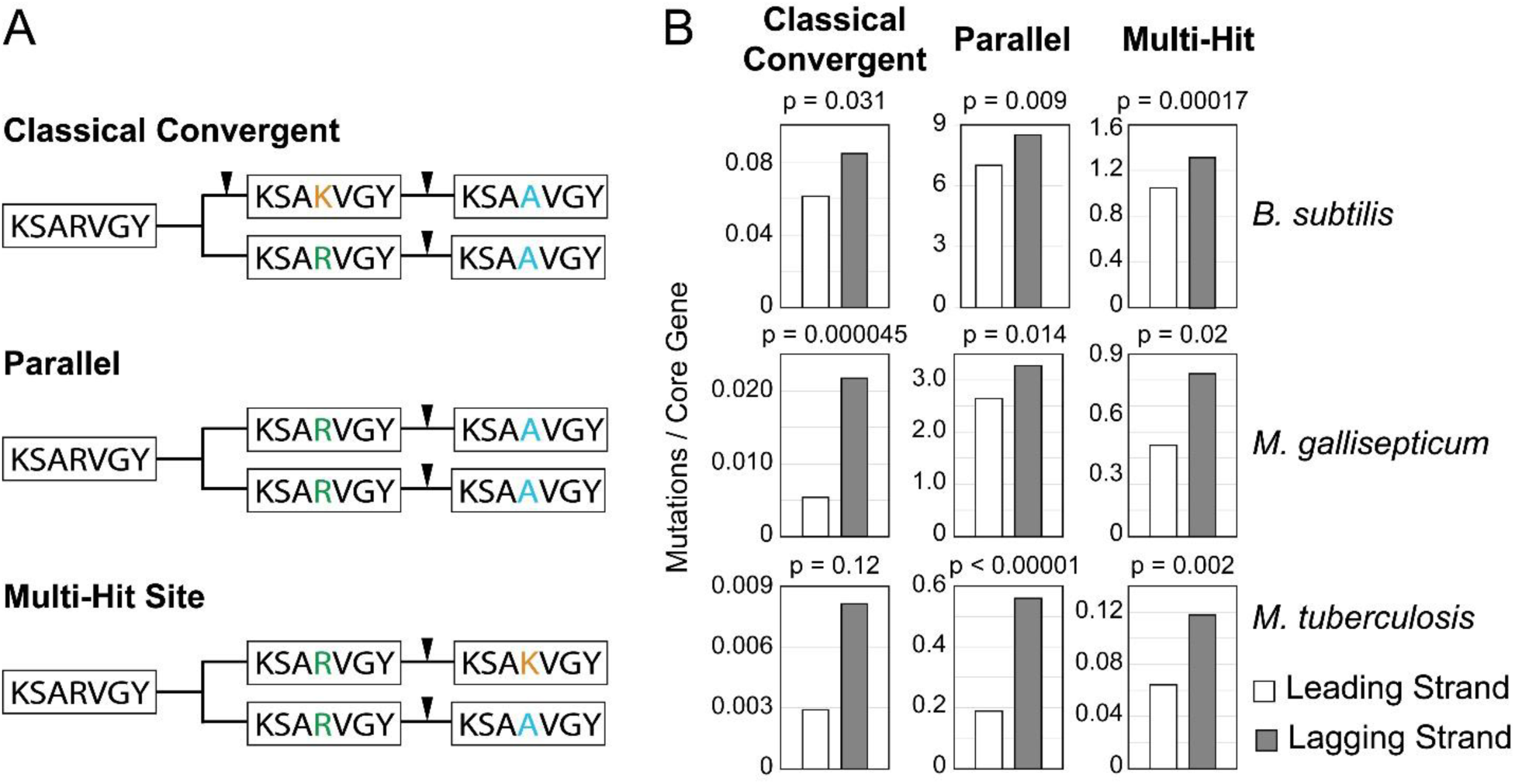
All three types of convergent mutations are more frequently observed in lagging strand genes. A) Three examples of convergent mutations. In each example, the amino acid sequence of a hypothetical gene fragment is shown for five species in a bifurcating lineage that evolves over time from left (older) to right (newer). Black arrows indicate the development of a nonsynonymous mutation in the displayed fragment of the example gene. B) The frequency of observed convergent mutations in either leading or lagging strand core genes of three species. The Chi square test was used to calculate significance.

To determine if lagging strand encoding can be adaptive, we conducted analyses of molecular convergence in three bacterial species *Bacillus subtilis, Mycoplasma gallisepticum*, or *Mycobacterium tuberculosis* using SNPs identified in natural isolates. We then identified the relative frequency of three classes of convergent mutations in leading and lagging strand genes: multi-hit site mutations, parallel mutations, and classical convergent mutations (Fig 1A). Below, our new analyses show that each type of convergent mutation is more common in lagging strand genes relative to leading strand genes. We further show that their abundance is both gene length and orientation-dependent, supporting our original model that lagging strand convergent mutations arose largely through mutagenesis caused by head-on replication-transcription conflicts. Our *in silico* modeling experiments further demonstrate that the observed mutations are highly unlikely to have arisen by chance alone, undermining the case for the neutral hypothesis. Our observation of the same patterns of convergence in three separate species strongly suggests that these trends are broadly conserved. As such, our results support the adaptive hypothesis. We conclude that lagging strand encoding can benefit cells through head-on conflict-mediated mutagenesis and the accelerated discovery of adaptive mutations.

## Methods

### SNP Analysis Using TimeZone

Accurate phylogenies are critical for identifying the independent emergence of same-site mutations^18,19^. As such, analyses of single bacterial species can be difficult due to the high degree of genetic similarity between individual isolates. For example, reports have shown that the commonly used codeml application of PAML can be limited in its ability to identify evidence of recent selection^18,20–22^. One solution to this problem is to develop zonal phylogenies in which nonsynonymous mutations are used to distinguish primary populations that differ only by synonymous mutations, from external populations that have undergone discrete non-synonymous changes^18^. Synonymous changes in the primary populations are interpreted as an indication of longer term stability, distinguishing these alleles from the more recently evolved external strains which each encode only non-synonymous changes^18^. This methodology, employed by the SNP analysis program TimeZone, facilitates the accurate identification of recently developed convergent mutations^23^. This program also reduces erroneous inferences of convergence by identifying and parsing out horizontally acquired mutations via MaxChi or PhylPro^24,25^. As such, TimeZone is optimized for identifying recent molecular convergence in closely related bacterial strains.

### Internal controls

The mutability hypothesis suggests that lagging strand genes, as a group, may tend to acquire more mutations than leading strand genes^11^. To avoid the possibility of comparing leading strand genes to potentially more mutable lagging strand genes, we restricted our analyses to core genes, defined as being 95% conserved in terms of amino acid content and gene length. This should normalize for any significant differences in purifying selection in either group. Our methods also include a second internal control: By comparing mutations in the leading versus lagging strand genes of the same isolates, any potential flaw in the inferred phylogenies generated by TimeZone should apply equally to leading and lagging strand genes. Therefore, erroneous data points should be equally abundant in both populations.

### Genomic Data

Bacterial genome files in were downloaded from NCBI in Genbank format and are listed in Table S1. Genomic sequences were analyzed using the program TimeZone version 1.0^23^. TimeZone parameters were adjusted such that only genes that are 95% conserved in both gene length and amino acid content were analyzed. Following TimeZone analysis, convergent mutations were parsed from using a custom script Analysis_conv5.py.

### Simulations

Simulations were performed as previously described, with one exception: all simulated multi-hit sites were mutated to a second amino acid^11^. As a result, a subset of our multi-hit site mutations were identical and thus represent parallel mutations. All code used for these simulations are publicly available at https://github.com/lh64/MultihitSimulation. As TimeZone outputs approximately 40k files per analysis, it was impractical to include this data in the supplement. However, all source data are available upon request.

## Results

To identify patterns of molecular convergence in leading or lagging strand genes, we used the program TimeZone to analyze point mutations (SNPs) in 50 fully assembled *B. subtilis* genomes. (See Methods for a discussion of internal controls and software settings.) We then parsed the convergent mutations identified by TimeZone into three groups: classical convergent, parallel, or multi-hit site mutations, and calculated their frequencies in leading or lagging strand core genes (Figure 1A, 1B, Methods). Our analysis identified a higher frequency of all three types of convergent mutations in lagging strand genes (Fig 1B). These results are highly statistically significant, supporting the hypothesis that positive selection acts more frequently on lagging strand genes.

To determine if our findings in *B. subtilis* are indicative of a broader pattern of evolution, we conducted the same analysis in a second Gram-positive species, *Mycoplasma gallisepticum*, and an unrelated species from a second phylum, *Mycobacterium tuberculosis* (Fig. 1C). We chose the *M. gallisepticum* because we previously found that it has the strongest correlation between gene orientation and mutation rates^6^. We selected *M. tuberculosis* as it is generally agreed that this species is unable to horizontally acquire DNA^26,27^. This was intended to address the formal possibility that our previous analyses could have been affected by recombination events. For each species, we observed the same trend as that identified in *B*. subtilis, strongly suggesting that the elevated frequency of convergent evolution is a conserved feature of lagging strand gene evolution.

### Most convergent mutations are retained due to positive selection, not chance

Classical convergent and parallel mutations are considered standard indicators of adaptation because they are extremely unlikely to occur by chance^11,28^. However, it is theoretically possible that multi-hit site mutations could arise at random with significant frequency^11^. Accordingly, they represent a lower confidence indicator of convergence. To determine if the homoplasy we observed would be expected under neutral conditions, we simulated a random distribution of the observed non-synonymous mutations in each gene as previously described^11^. Specifically, we drew random variable sites within each core leading or lagging strand gene with replacement until the total number of sites equaled the number of observed amino acid changes^11^. For any site drawn twice, the original residue was randomly mutated to one of the 19 other amino acids, yielding either a multi-hit (different mutations) or parallel mutation (identical mutations). We performed simulations for all leading and lagging strand core genes using nonsynonymous substitution data from either our original 5 genome or new 50 genome study of *B. subtilis*. We repeated these simulations 10,000 times for all leading or lagging strand core genes, yielding a distribution of values (Fig. 2). We found that the observed number of both parallel and multi-hit site mutations are greatly in excess of even the most extreme simulated data (Fig. 2). Therefore, the data strongly suggest that the multi-hit site and parallel mutations we observed in both our original and current studies did not arise by chance (Fig. 2). Instead, they most likely arose through positive selection, consistent with the idea that both multi-hit site and parallel mutations are indicative of evolutionary convergence.

**Figure 2.**
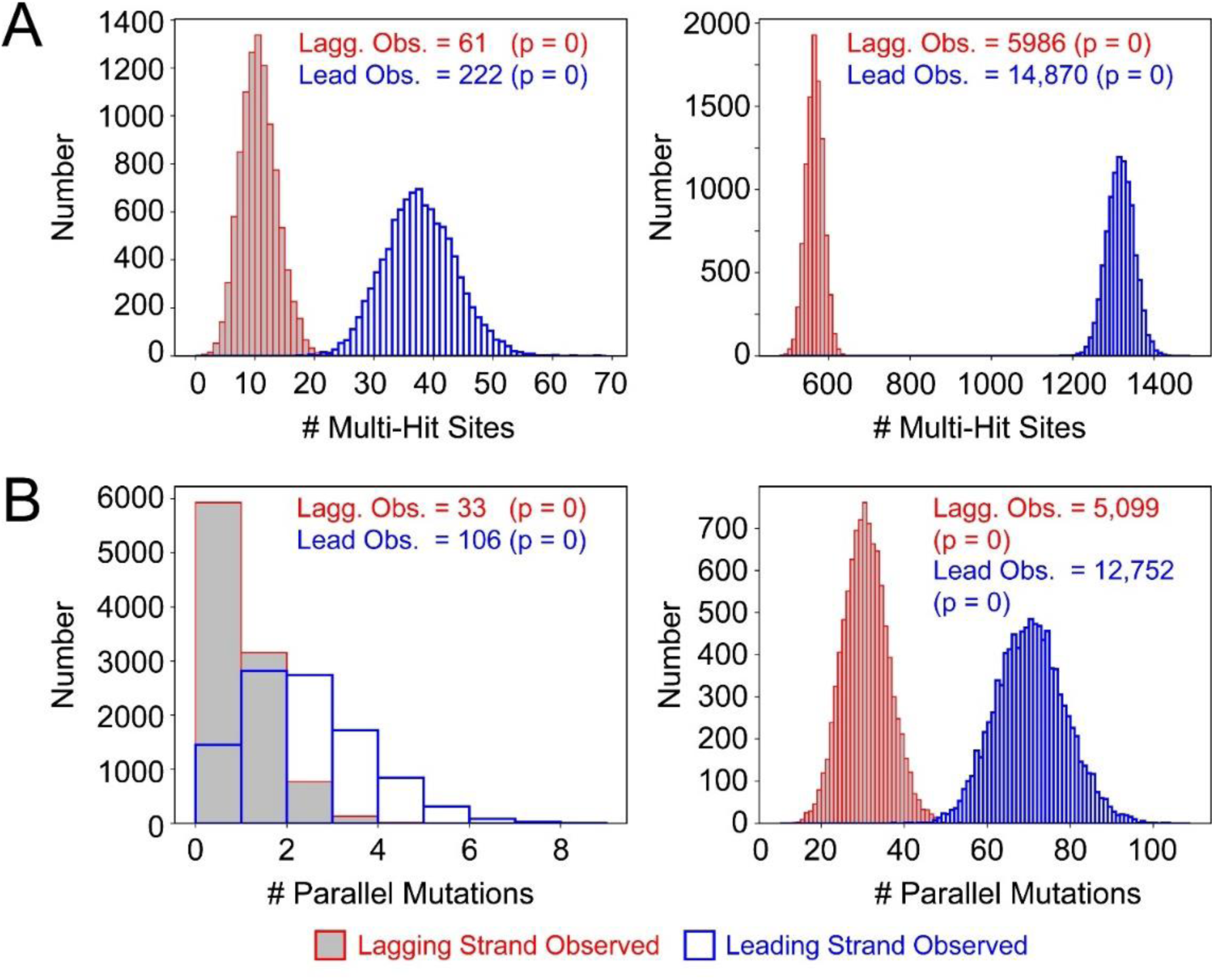
Observed convergent mutation frequencies cannot be explained by chance. A) The distribution of multi-hit site mutations expected under neutral selection conditions was determined by simulating a random reassortment of the observed nonsynonymous substitutions in leading strand (blue) or lagging strand genes (red) based upon the numbers identified in either our previous 5 genome analysis of *B. subtilis* (left graph), or our 50 genome analysis (right graph). This was performed for all leading or lagging strand genes x 10,000 iterations, providing a distribution for both groups. The actual observed numbers are shown above the simulated distributions. B) Lower graphs: Same as above, but for parallel mutations.

### Head-on replication-transcription conflicts increase the frequency of convergent mutations in lagging strand genes

Our previous work indicated that head-on replication-transcription conflicts are the mechanistic basis for the increased mutation rate of head-on genes^6,7,29^. Evidence for this hypothesis includes both experimental results, and our observation of a gene length and orientation-dependent increase in both dN/dS ratios and mutation frequency^7^. This result was consistent with the idea that head-on conflict severity should increase in direct relation to gene length, whereas co-directional conflicts (leading strand genes) should not^7,30^. If head-on replication-transcription conflicts are responsible for promoting the formation of the observed convergent mutations identified here, their abundance should follow the same pattern. To test this, we calculated the number of convergent mutations per core gene, then assessed the relationship with gene length (Fig. 3). We found that all three types of convergent mutations increase in frequency in a gene length-dependent manner. We also found that this effect is more pronounced in lagging strand genes, strongly suggesting that head-on conflicts are indeed responsible for the increased frequency of convergent mutations in lagging strand genes.

**Figure 3.**
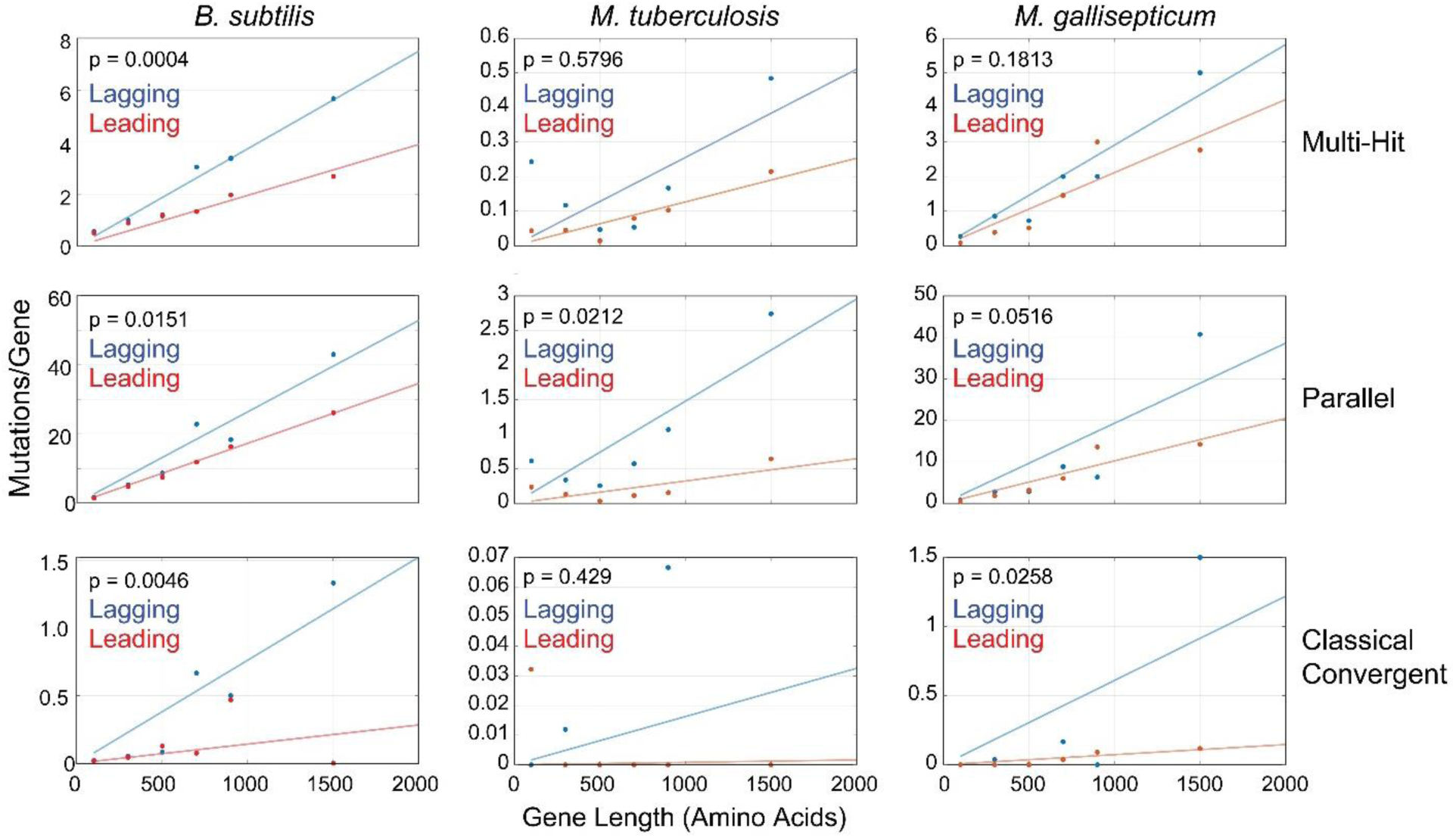
The frequency of convergent mutations increases in a gene length and orientation-dependent manner. The number of convergent was calculated for genes of varying length. The convergent mutation type is indicated to the right of each row of graphs. Genes were binned into 6 groups based upon their length in amino acids: 1) <200 AA, 2) 201-400 AA, 3) 401-600 AA, 4) 601-800 AA, 5) 801-1000 AA, 6) 1001-2000 AA. F-test p-values are indicated in the top left.

To determine if this mechanism is conserved, we performed equivalent mutational analyses in two additional species: *M. gallisepticum* and *M. tuberculosis*. In both cases, we identified the same convergent mutation pattern found in *B. subtilis* (Fig. 3).

## Discussion

The new analyses presented here demonstrate that lagging strand genes accumulate all three types of convergent mutations at a higher rate than leading strand genes, and that this effect is broadly conserved. Our modeling experiments indicated that both parallel and multi-hit site mutations are unlikely to have arisen by chance alone, undermining the case for the neutral hypothesis. As such, our findings strongly support the idea that the elevated mutation rate of lagging strand genes is due to positive selection rather than higher mutability.

The evidence for positive selection presented here is consistent with the results of our previous study in which we identified a higher frequency of genes with a dN/dS ratio significantly greater than 1 encoded on the lagging strand of six diverse bacterial species^6^. This metric represents a second independent indicator of positive selection, and cannot be explained by relaxed purifying selection^31^. Therefore, both data sets directly support the adaptive hypothesis for lagging strand encoding.

Our simulations were intended to test the possibility that neutral evolution could explain the higher frequency of convergent mutations on the lagging strand. However, rather than a completely neutral simulation, we performed a semi-neutral simulation based the observed non-synonymous mutation rates, which, by definition, are the result of both mutagenesis and selection^11^. As the dN is significantly higher for lagging strand genes, the simulations provided more chances for these genes to gain multi-hit site or parallel mutations by chance alone. Therefore, our setup represent a more conservative approach than a fully neutral simulation based upon mutation rates (dS values) which are equal for both groups^6^. As a result, our simulated distributions probably overestimate the frequency of non-adaptive convergent mutations in lagging strand genes. In spite of this, our simulations still indicate with high confidence (p = 0 in all cases) that the neutral model is not sufficient to explain the vast majority of convergent mutations in either leading or lagging strand genes.

The overestimated rate of non-adaptive convergence in lagging strand genes may explain why a previous group that conducted a roughly equivalent simulation came to the opposing conclusion^11^. Noting that the observed ratio (lagging/leading) of convergent mutations matches the simulated ratio, they inferred that the neutral model is correct^11^.

However, the ratio of simulated values is unlikely to be an informative metric for two reasons: 1) Being based on the dN, the simulated lagging/leading ratio is probably an overestimate, and 2) chance can only explain, for example, 0.5-1% of the observed parallel mutations (Figure 2B, right panel). Therefore, for the neutral model to be correct, the rate of non-adaptive (chance) convergence would need to be approximately 100 to 200-fold higher than the simulated rates which is highly unlikely.

It is also important to note that the equal dS of leading and lagging strand genes does not contradict the idea that, when transcribed, lagging strand genes have a higher mutation rate^7,29,32^. As the dS is an average, it accounts for the frequency and variety of mutation rates during both transcriptional induction (high mutation rate) and repression (low mutation rate)^7,29,32^. Our previous work suggests that lagging strand is enriched in various types of stress response genes which are conditionally induced, whereas the leading strand preferentially encodes constitutively and highly transcribed genes^5^. Therefore, the equal dS may simply reflect the less-frequent transcriptional induction of lagging strand genes despite their higher mutation rate when transcribed.

Our model suggests that, in addition to positive selection, other evolutionary pressures act in parallel on lagging strand genes. For example, other studies have emphasized the idea that head-on replication-transcription conflicts should confer negative selection against highly transcribed lagging strand alleles^3,11,15,33^. This mutation-selection balance hypothesis has been presented as a competing and mutually exclusive explanation to the adaptive hypothesis^11,15^. Though we agree that the mutation-selection balance may have an important influence on lagging strand genes, we disagree that the two models are mutually exclusive. The case for positive selection is simply too strong to reasonably ignore. Of course, it is unlikely that all lagging strand genes are under positive selection, leaving room for other influences. Therefore, we propose a unified model in which a combination of positive and negative selection, as well as neutral evolution, collectively drive the retention of genes on the lagging strand.

## Acknowledgements

H.M. and C.M. were supported by the National Institute of Health, R01-AI-127422.

---

**Table S1.**
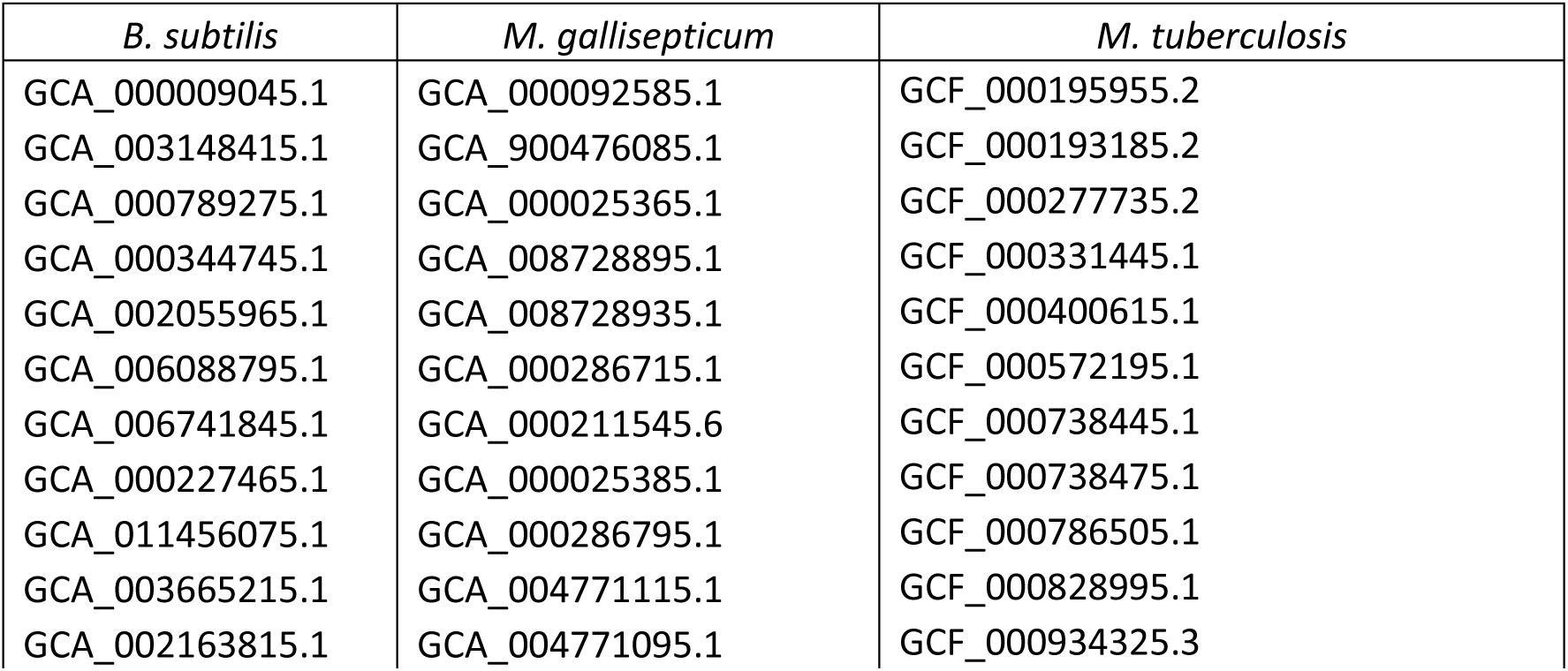
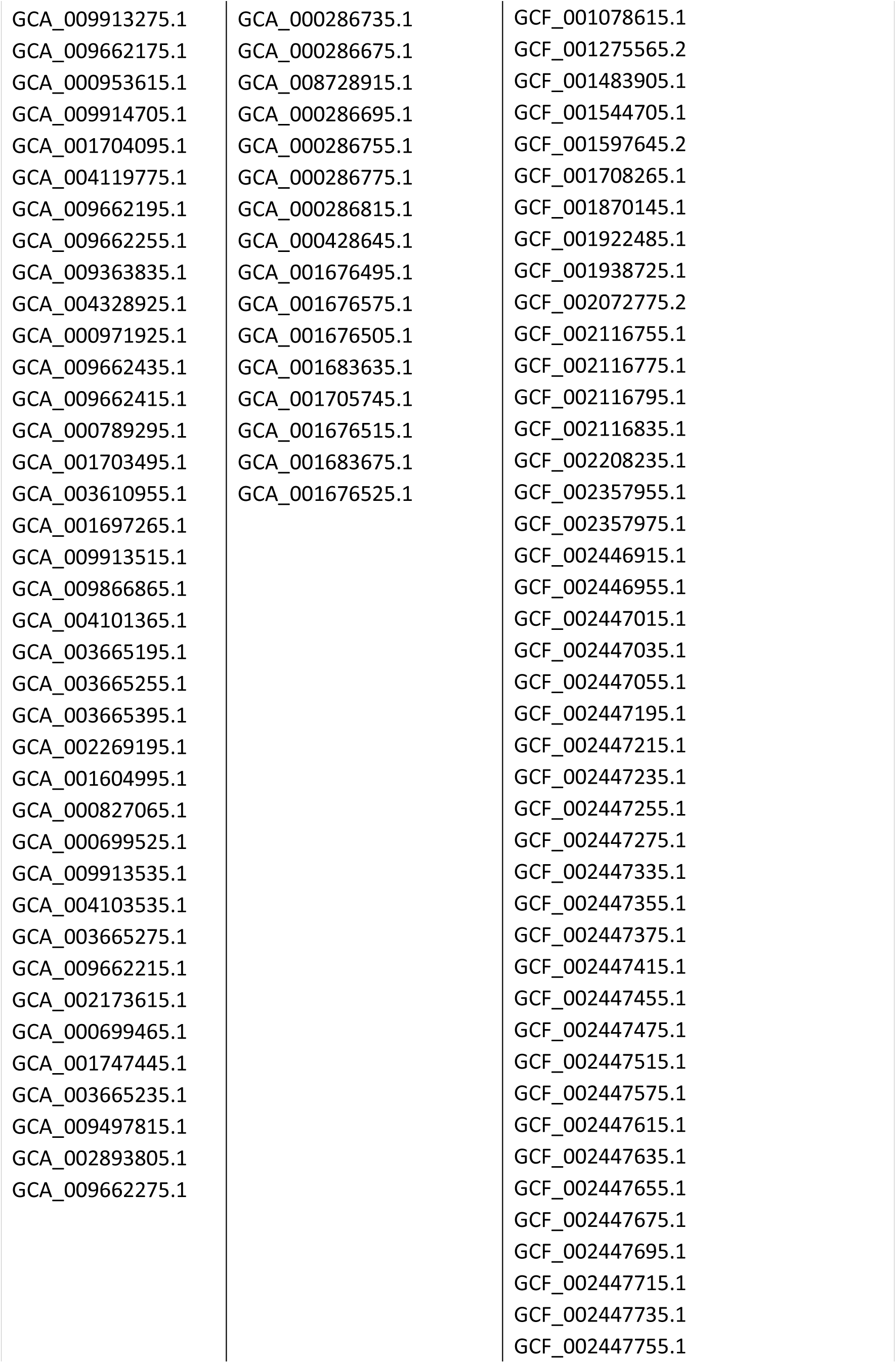
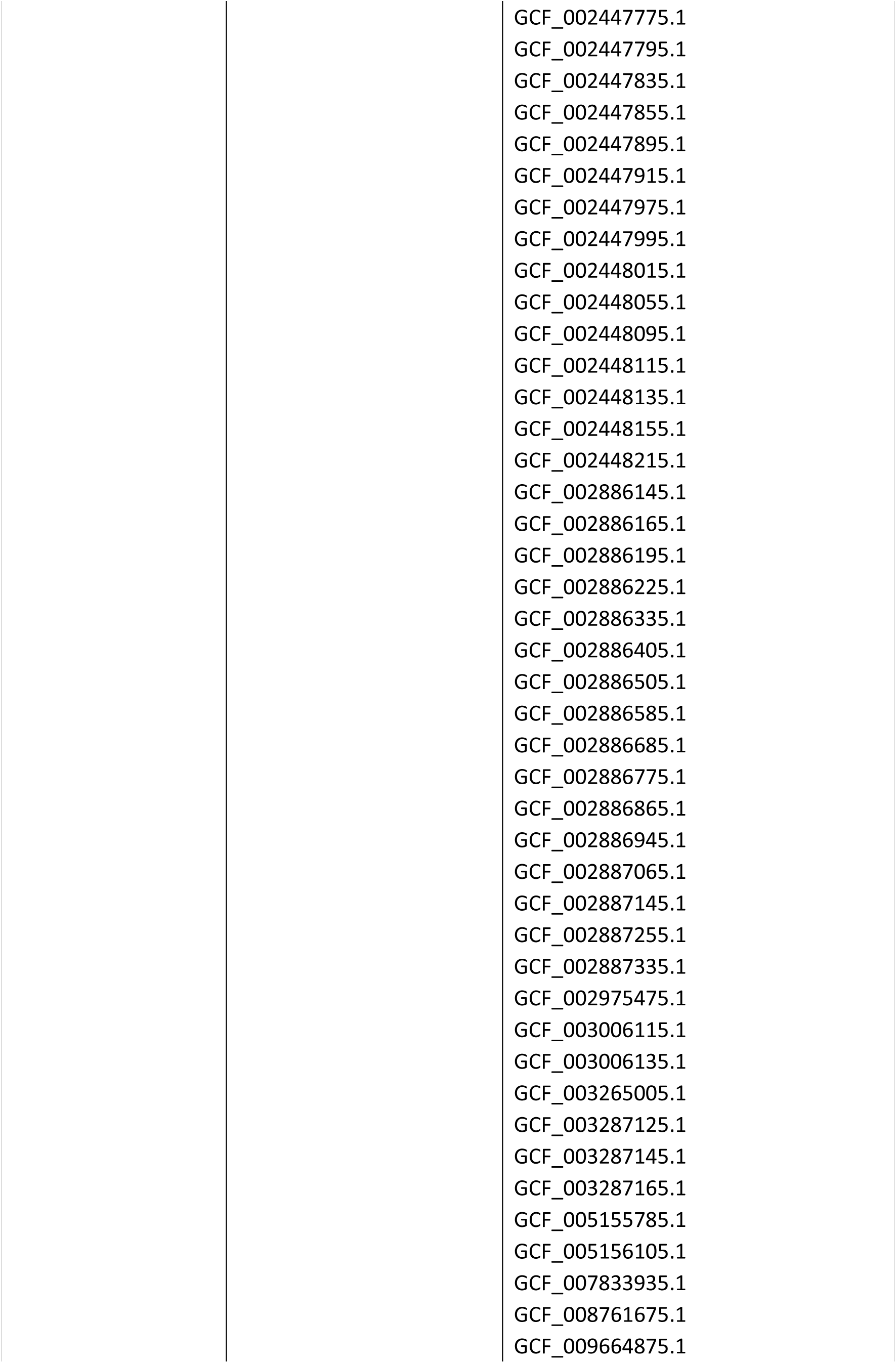
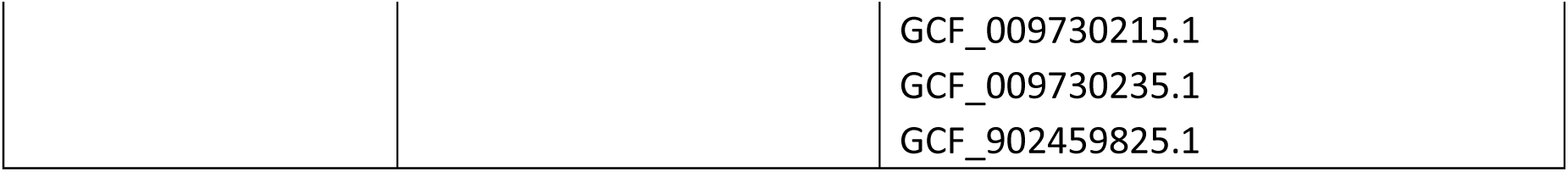
Genomes used for mutational analyses. The NCBI Genome assembly serial numbers for all genomes analyzed by TimeZone are listed. The Genbank formatted genomes were downloaded and used as input.

